# Mesostats — A multiplexed, low-cost, do-it-yourself continuous culturing system for experimental evolution of mesocosms

**DOI:** 10.1101/2022.04.20.488910

**Authors:** Erika M Hansson, Dylan Z Childs, Andrew P Beckerman

## Abstract

Microbial experimental evolution allows studying evolutionary dynamics in action and testing theory predictions in the lab. Experimental evolution in chemostats (i.e. continuous flow through cultures) has recently gained increased interest as it allows tighter control of selective pressures compared to static batch cultures, with a growing number of efforts to develop systems that are easier and cheaper to construct. This protocol describes the design and construction of a multiplexed chemostat array (dubbed “mesostats”) designed for cultivation of algae in 16 concurrent populations, specifically intended for studying adaptation to herbicides. We also present control data from several experiments run on the system to show replicability, data illustrating the effects of common issues like leaks, contamination and clumps, and outline possible modifications and adaptations of the system for future research.

## INTRODUCTION

Microorganisms provide an unparalleled opportunity for the study of evolutionary dynamics due to their combination of short generation time, simple genetics and ability to fit huge population sizes in a small space. The path of evolutionary adaptation can thus be replicated and tightly controlled in real time in the lab, allowing exciting new insights into the mechanisms of adaptive evolution and testing of predictions from theory (Barrick & Lenski, 2013; Good *et al*., 2017; Kawecki *et al*., 2012; Lang & Desai, 2014; Van den Bergh *et al*., 2018).

The most common way of growing microorganisms for experimental evolution is as batch cultures. This involves serial repetition of small cell population subsets being moved to fresh medium and grown to stationary phase before being transferred again to fresh medium to allow new growth and, with time, adaptation. This is a simple, cheap and scalable method, but its drawback is the resulting fluctuating environment as the cells go through “boom and bust”-cycles at every transfer resulting in a complex selective environment (Gresham *et al*., 2008; Gresham & Dunham, 2014; Van den Bergh *et al*., 2018). As the nutrients in the medium gradually run out, the cells will arrest growth and division while waste products build up. Oxygenation, light levels and pH will also fluctuate with population density. All of this affects cellular metabolism and physiology and subsamples taken from such populations will be growth phase specific, making it difficult to define and isolate the selective pressures acting on the populations (Gresham & Dunham, 2014). Furthermore, there is an evolutionary bottleneck at each transfer, where the considerable reduction in population size associated with transfer to the next batch affects the genetic diversity and mutational space available (Lenski *et al*., 1991; Van den Bergh *et al*., 2018; Vogwill *et al*., 2012; Wahl & Gerrish, 2001), giving an increased role to genetic drift in the evolutionary outcome (Elena & Lenski, 2003).

Chemostats – continuous flow-through, chemically stable cultures where growth medium and treatments are fed into the fixed-volume populations at a constant rate – solve these issues as the specific growth rate of the population at steady state is matched to the dilution rate (Monod, 1950; Novick & Szilard, 1950). The populations are maintained in exponential growth and constant mixing ensures a homogeneous environment, allowing precise control of the relevant selective pressures compared to the complex dynamics present in batch cultures (Gresham & Dunham, 2014). This unique opportunity for experimental manipulation offers a high-throughput chance to pick apart evolution in action and, as a result, chemostats have recently seen a renaissance in experimental evolution and systems biology as new technological advancements make them easier to maintain than ever before (reviewed in Bull, 2010; Gresham & Hong, 2014). Chemostats also allow following population fluctuations and evolutionary dynamics in response to experimental treatments in the long term, where equilibria and population cycles including several species and strains can be described as a function of the flow rate (e.g. Becks *et al*., 2012; Declerck *et al*., 2015; Fussmann *et al*., 2000; Hiltunen *et al*., 2014; Yoshida *et al*., 2003). Multiplexed arrays, where the dilution rate is set by a single pump, and medium sources can be shared, further minimise variation between population chambers (Dénervaud *et al*., 2013; Ekkers *et al*., 2020; Miller *et al*., 2013; Skelding *et al*., 2018; Tonoyan *et al*., 2020; Toprak *et al*., 2013; Wong *et al*., 2018).

Here we describe a multiplexed small-scale DIY chemostat array system (dubbed “mesostats”) adapted from the ministat array developed by Miller *et al*. (2013) to suit experimental evolution of algae, in contrast to the so far described designs specifically intended for yeast (Dénervaud *et al*., 2013; Miller *et al*., 2013; Wong *et al*., 2018) and bacterial cultures (Tonoyan *et al*., 2020; Toprak *et al*., 2013). Our system uses common algal model species *Chlamydomonas reinhardtii*, with the specific goal to use it as a herbicide resistance evolution model. *C. reinhardtii* is an established model species for herbicide resistance evolution (Lagator *et al*., 2013a,b; Reboud *et al*., 2007; Vogwill *et al*., 2012) and molecular analysis of herbicide resistance mutations (Erickson *et al*., 1984, 1989; Randolph-Anderson *et al*., 1998), but all studies to date have used batch cultures. We present the full protocol for assembling and maintaining a 16-chamber mesostat array by a single person as well as control data illustrating the ability of the system to track trends and variability in the abundance of organisms among replicates. We also present pilot data illustrating the ability to use the mesostats to evolve resistance in *C. reinhardtii* to growth inhibiting herbicide glyphosate. Furthermore, we have included data from this system illustrating the signal of common problems like leaks, contamination and cell clumping, showing how to distinguish it from biological variation as well as how to prevent and address these problems if they occur. We also outline the ways in which this system could be further modified and avenues of future research.

## METHODS

### AN OVERVIEW OF THE DESIGN

The mesostat array consists of four main parts: (1) the medium line, (2) the culture chambers, (3) the overflow chambers and the (4) aeration line. Medium is pumped from the medium containers via a peristaltic pump into the culture chambers (mesocosms) where the experimental organisms are grown. Air is pumped through a gas washing bottle into the culture chambers via the aeration needle to ensure mixing and create pressure so liquid flows out through the efflux needle and the culture stays a fixed volume. The efflux line leads to an overflow chamber, and the volume collected in these is regularly measured to ensure equal flow rates between all culture chambers. Samples for analysis can be obtained from the overflow, but this only samples from the top of the culture and the environment in the collection receptacle may differ from the culture chambers. To allow sampling from lower levels of the culture, sampling needles have been fitted to the culture chambers. The sampling needle can also be used for inoculation or addition of treatments to the chambers. The entire setup can be kept in a controlled temperature room which ensures low levels of evaporation, but the culture temperature can also be maintained by other methods such as a light table or a water bath. When growing photosynthetic organisms such as algae, even light levels for all chambers are best maintained by a light table as well as fitting strip lights around the chambers. An overview of the full design is seen in Figure 1 and Figure 2A, as well as full assembly schematics in item Figure 3–5. The control conditions are summarised in Table 1.

**Figure 1:**
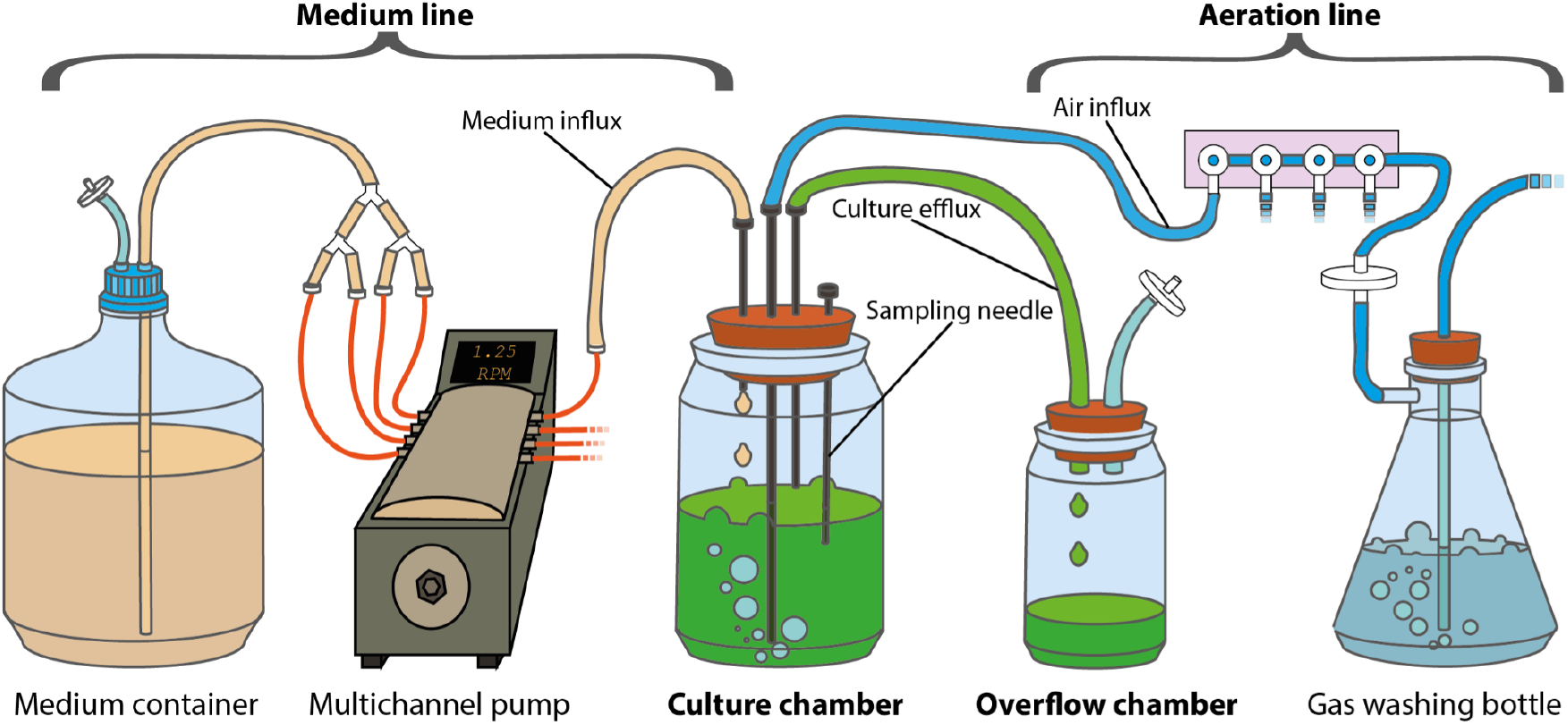
Simplified schematic overview of the mesostat system showing medium containers, the pump, the culture chamber with sampling needle, the overflow bottle, the gas washing bottle along with the medium and air influx lines and the culture efflux line.

**Figure 2:**
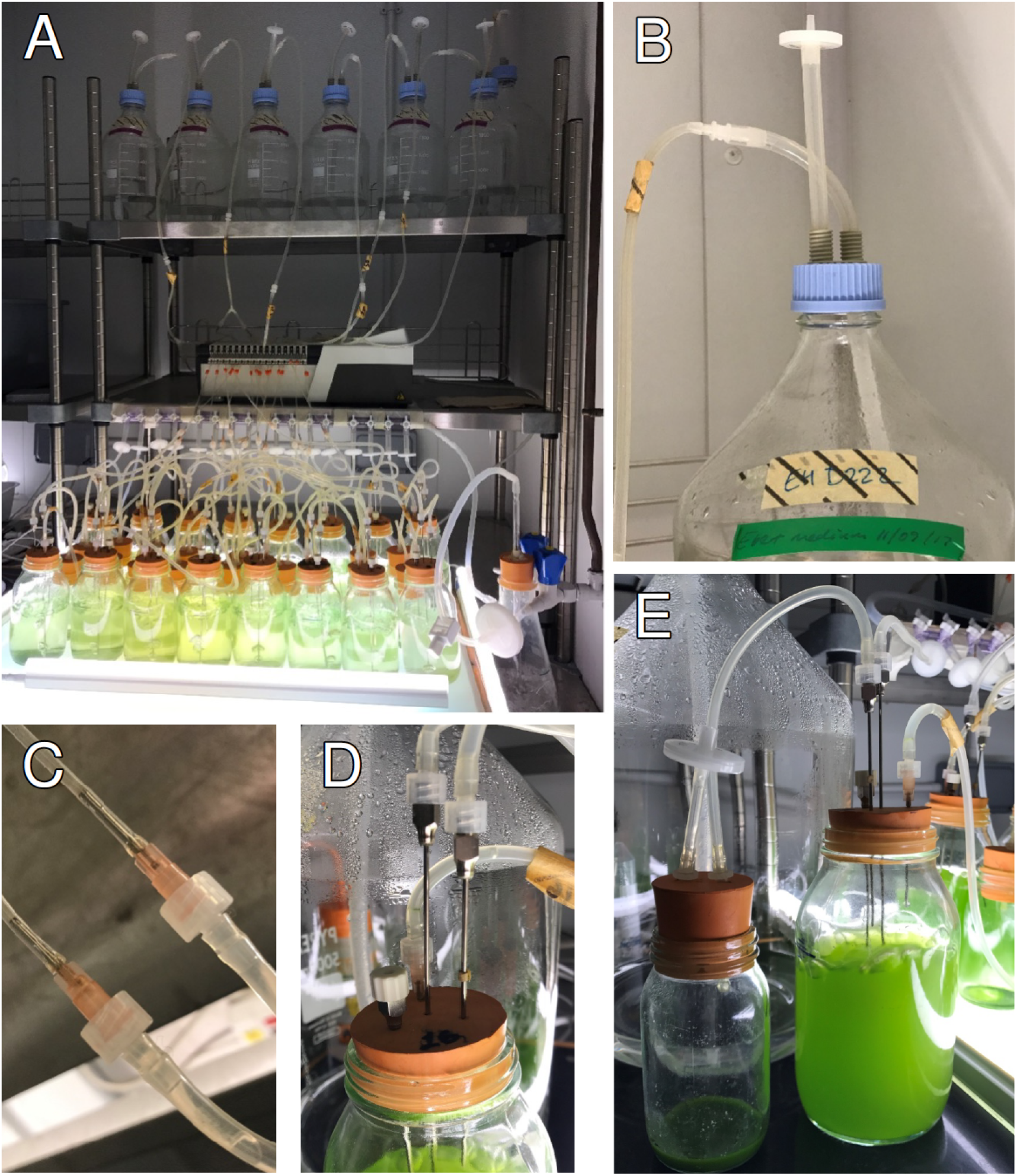
A) The complete setup just after inoculation with algae, running an experiment with six levels of treatments applied through the media lines. B) Close-up of medium siphon through medium container lid. C) Close-up of connection between pump tubing and silicone tubing used throughout array. D) Close-up of culture chambers rubber bung with the four hypodermic needles, capped sampling needle to the left in foreground, steel efflux needle to the right in foreground, steel aeration needle in the middle, and pink plastic medium influx needle in the background. E)The overflow chamber (left) and the culture chamber at steady state (right) with the efflux line running between them.

**Figure 3:**
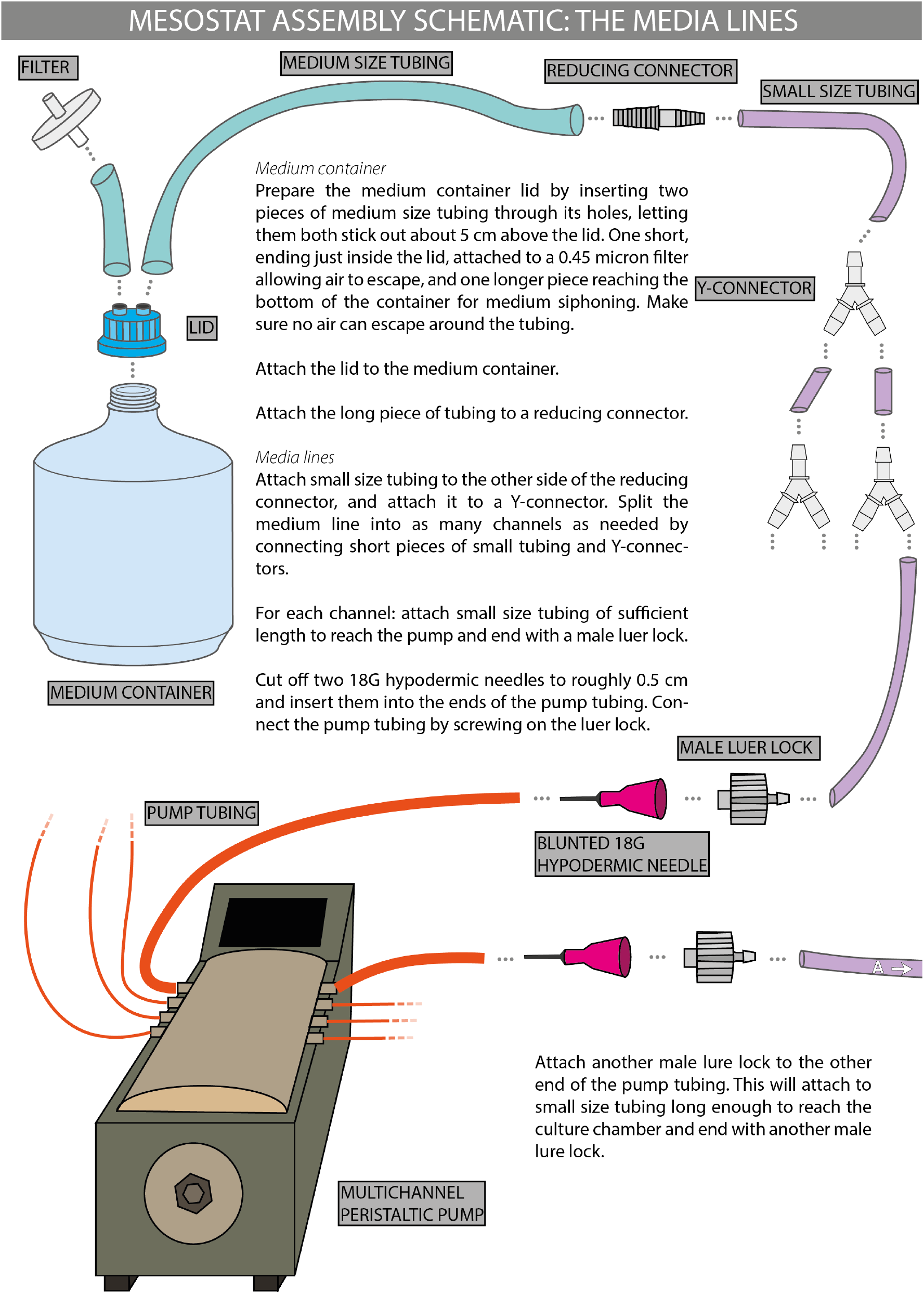
Assembly schematic part 1, the media lines.

**Figure 4:**
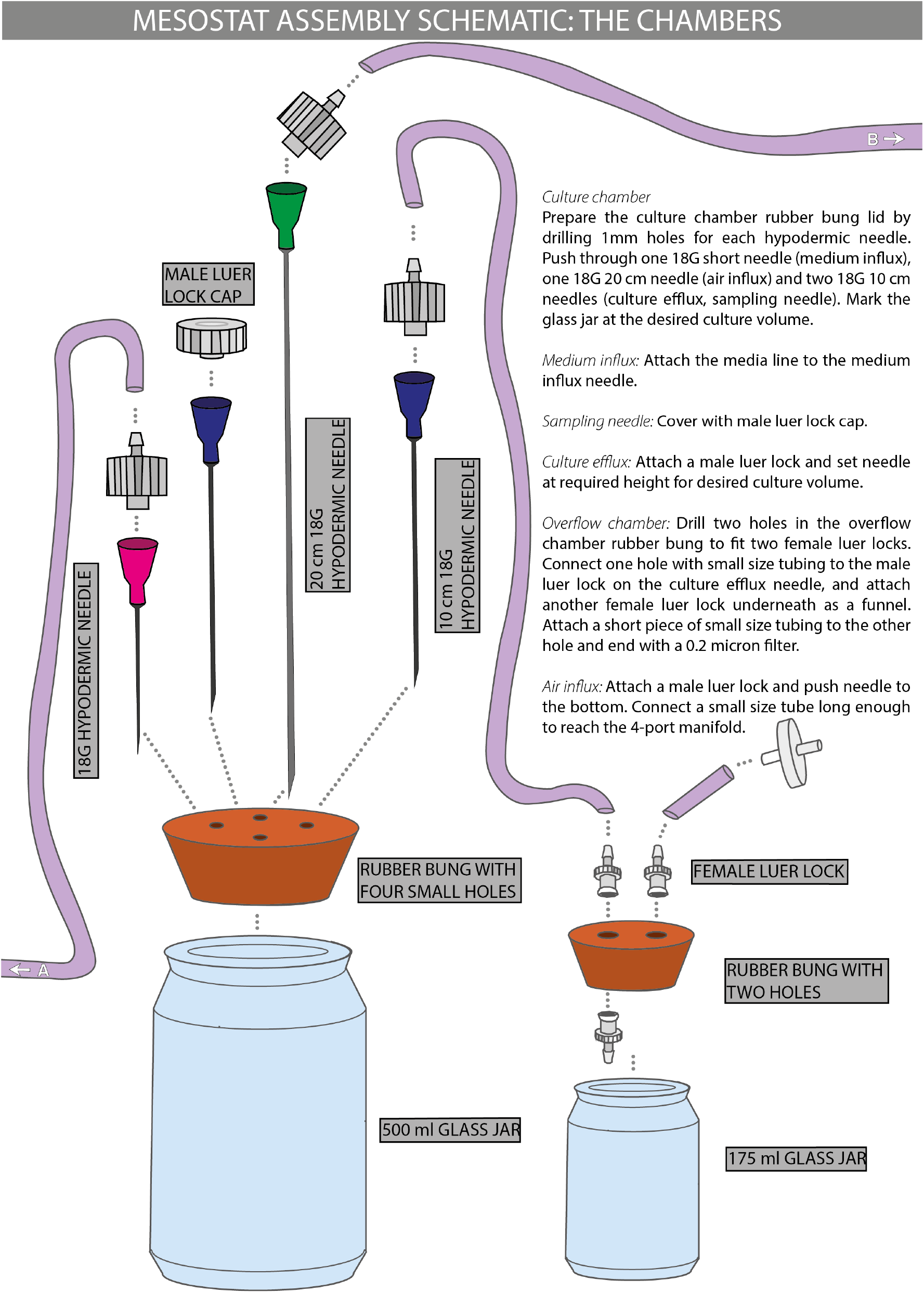
Assembly schematic part 2, the chambers.

**Figure 5:**
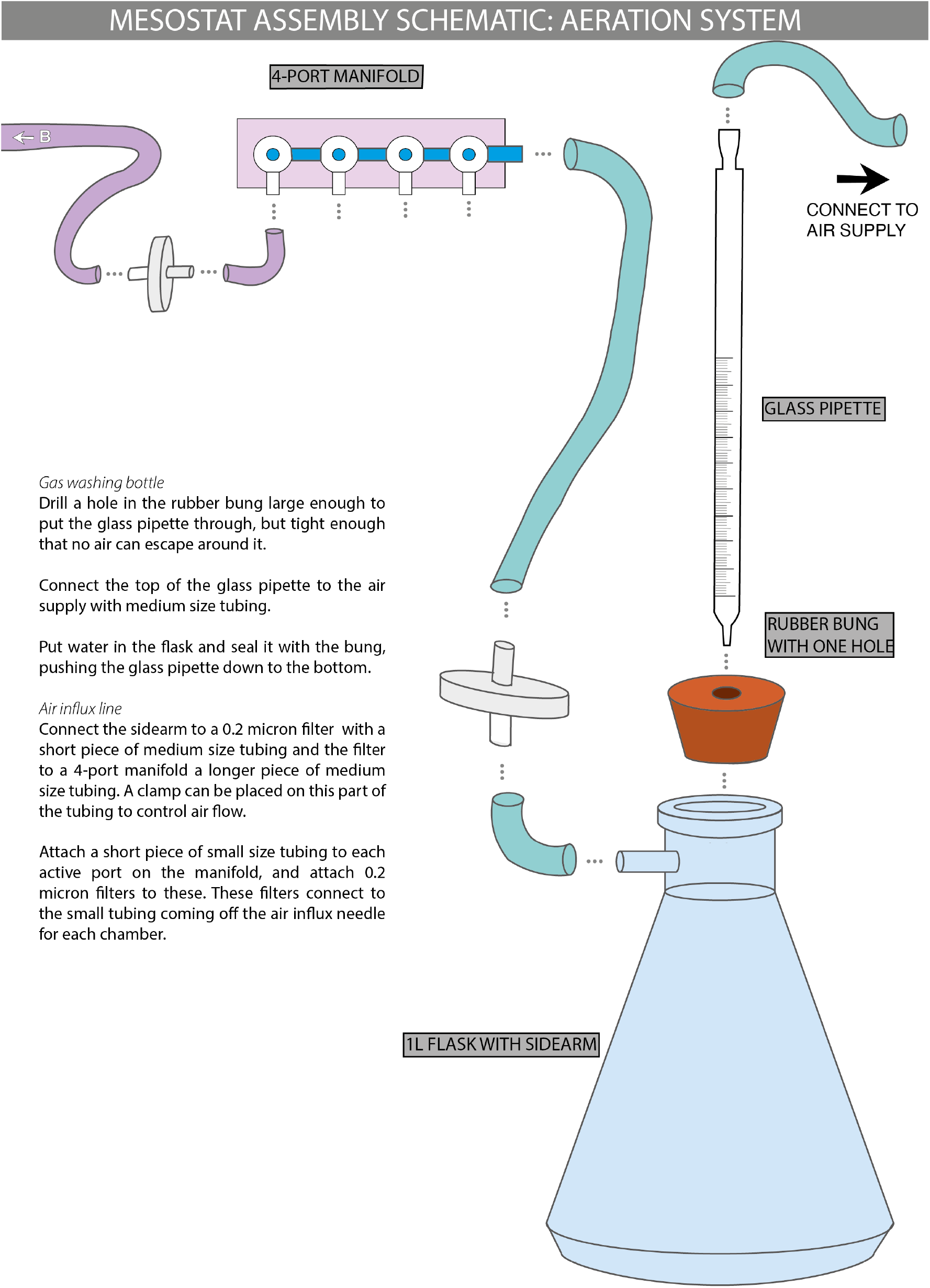
Assembly schematic part 3, the aeration system.

**Table 1:** Control conditions for *C. reinhardtii* cultures.

|  |  |
| --- | --- |
| **Light level** | 75 $\mu\text{mol m}^{-2} \text{s}^{-1}$, 24 h, all directions |
| **Internal culture temperature** | 30°C |
| **CT room ambient temperature** | 25°C |
| **Medium flow rate** | 0.15/day, 1.25 RPM pump speed |
| **Sampling frequency and volume** | 1.5 ml day |
| **Medium** | Ebert algal medium ((Ebert, 2013)) |

#### THE MEDIUM CONTAINERS

The volume and number of medium containers depends on the experimental design and number of treatments. When the experimental design calls for different treatments applied via the medium, culture chambers sharing a treatment share a medium source. The volume of the medium containers should be chosen to allow sufficient medium to supply all its chambers for at least 5 days, to ensure the time to prepare new sterile medium before changing is needed. The depletion rate of the medium will depend on the number of chambers sharing a container and the flow rate.

The medium container should be sealed with a lid or stopper with a hole that allows air to escape through a filter. If the medium outflow can be through either a tap at the bottom of the container, or tubing through the lid siphoning medium from the bottom.

For the data presented in this thesis, a single 20 L autoclaveable glass container, fitted with a silicone stopper and a filter (0.45 μm) on top and a tap connected to large silicone tubing (inner/wall diameter: 12.5/2.25 mm) at the bottom, is used when all 16 chambers are receiving the same medium.

For data obtained from experiments where different treatment levels were present, 5 L or 2 L autoclaveable glass bottles are used for the treatments, each fitted with medium silicone tubing (inner/wall diameter: 6.5/1.5 mm) siphoning the medium out through holes in the lids alongside tubing ending in a filter to allow air to escape the bottle (Figure 2B). For example, as seen in Figure 2A, 6 different herbicide concentrations are applied through the medium to two chambers each from 2L bottles, and the control treatment is supplied to four chambers from a 5L bottle.

#### THE PUMP AND THE MEDIUM LINES

The medium is pumped from its container(s) by a multichannel peristaltic pump. Our design employed a Watson-Marlow 205S/CA16 (16 channels) and pumped liquid through small silicone tubing (inner/wall diameter: 3/1 mm), connected to the medium container tubing with a reducing connector. The tubing is split using Y-connectors before the pump so that each culture chamber has its own media line. The same tubing is used throughout the mesostat array (Figure 2A), except for the tubing mounted in the pump itself which is autoclaveable marprene tubing (Watson-Marlow, orange/orange, 0.88 mm bore), connected to the silicone tubing using cut off and blunted hypodermic needles (18 G, 50 mm) inserted into the pump tubing and connected to the silicone tubing using male luer locks (Figure 2C).

#### THE CULTURE CHAMBERS

Each culture chamber consists of a 500 ml glass jar, sealed with a rubber bung. The rubber bung has four hypodermic needles inserted through it (Figure 2D): a medium influx needle (18 G, 50 mm) connected to the tubing running through the pump, an aeration needle (16G, 203 mm) connected to the aeration system allowing constant mixing of the culture, a sampling needle (16 G, 203 or 101 mm depending on sampling needs), and an efflux needle (16 G, 101 mm) which sets the culture volume.

Tubing is connected to the medium influx, aeration, and efflux needles using male luer locks. A sterile syringe is used with the sampling needle to pull samples out of the chamber or for injections, which is kept sealed with a male luer cap when not in use.

#### THE AERATION SYSTEM

The culture chambers have a constant influx of air for mixing to prevent the organisms – in our case algae – from sedimenting, and to create pressure for the efflux of liquid so that the culture is kept at a constant volume. Air can be supplied by lab/building infrastructure, e.g. air supply taps, or by an aquarium pump with high enough pressure. The air passes through a filter (0.2 μm) into a gas washing bottle to prevent evaporation. The gas washing bottle consists of a 1L flask with a sidearm, with the incoming air being passed through a glass pipette into distilled water. The air is pushed from the gas washing bottle through 4-port manifold connectors (180° rotation), splitting the air supply to tubing for each culture chamber and passing it through a second filter (0.45 μm). This tubing connects to the aeration needle fitted in the culture chamber bung. The air supply can be controlled using an adjustable clamp fitted to the tubing between the gas washing bottle and the manifold connector.

#### THE OVERFLOW CHAMBERS AND THE EFFLUX LINE

The efflux line from each culture chamber leads to an overflow chamber consisting of a 175 ml glass bottle sealed with a rubber bung (Figure 2E). The efflux tubing is connected to a hole in the rubber tubing using a female luer lock. A second female luer lock is also fitted on the opposite side of the hole on the underside of the bung, to serve as a funnel for the incoming liquid. A second hole connected with a female luer lock to tubing ending with a filter (0.45 μm) allows air to escape the collection chamber. This chamber should be emptied regularly and can be used to control that the flow rate remains equal between the culture chambers. Samples can be obtained from the overflow chamber, but the environment in the overflow chamber will be different from the culture chambers. With a low flow rate and a high temperature environment, the culture will evaporate quickly when its volume is low. The overflow chamber is also not being diluted with fresh medium, meaning the cells are no longer kept in exponential growth or connected to the aeration line, often resulting in sedimentation and stratification of the culture.

### THE LIGHT SYSTEM

The light is provided by white light LED strip lights mounted around the chambers and between the two rows of chambers, as well as a DIY light box consisting of white light LED strip lights and a semi-transparent plastic top to diffuse the light. Equal light from all angles is essential to ensure even algal growth in the chambers. A light box is not necessary, but convenient and can be used for providing light to batch cultures or growth assays of subsamples.

### MATERIALS AND EQUIPMENT

Here we present a complete list of materials required to construct a 16-chamber array. Note that the media containers and associated lids are listed as optional, as exact size and number needed depend on the experimental design. If the company and product code is not listed, the part was not acquired new and the exact same product is no longer sold. Other than the pump and pump tubing, all of the pieces are fairly standard pieces found in many wet labs and similar products can be obtained easily from all major scientific suppliers.

1. Small tubing, 20 m (Fisherbrand™ Silicone Tubes; inner/wall diameter: 3.0/1.0 mm, Fisher Scientific, 10111801)
2. Medium tubing, 20 m (Fisherbrand™ Silicone Tubes; inner/wall diameter: 6.5/1.5 mm, Fisher Scientific, 10549201)
3. Large tubing, 10 m (Fisherbrand™ Silicone Tubes; inner/wall diameter: 12.5/2.25 mm, Fisher Scientific, 10726931)
4. Reducing connector, 10 (Reducing Connector PVDF; 1/4” to 1/8”, Cole Parmer, EW-30703-50)
5. Male luer lock, 100 (Cole-Parmer ADCF Male Luer to 1/8” L Barb Adapter, Cole Parmer, WZ-30800-24)
6. Female luer lock, 50 (Cole-Parmer ADCF Female Luer to 1/8” L Barb Adapter, Cole Parmer, WZ-30800-08)
7. Y-connector, 40 (Barbed Y Connector; PVDF; 1/8”, Cole Parmer, WZ-30633-44)
8. Straight connector, 10 (Barbed fittings; Straight Connector; Kynar; 1/4” ID Cole Parmer, WZ-30703-05)
9. Gas washing bottle, 1 (1L flask with sidearm tubulation)
10. Glass pipette, 1 (10 ml glass pipette)
11. 4-port Manifold, 4 (Polycarbonate individual manifolds with luer locks; 4 ports; 180° rotation Cole Parmer, EW-06464-85)
12. 0.45 μm filters, 100 (PTFE Nonsterile Syringe Filters; 0.45 micron; 25 mm dia, Cole Parmer, WZ-02915-22)
13. 0.2 μm filters, 1 (AcroVent 0.2μm PTFE, Pall Corporation, 4249)
14. Air tubing clamp, 1 (Adjustable tubing clamp)
15. Multiplexed peristaltic pump, 1 (205S/CA16 16 Cartridge pump, Watson-Marlow, 020.3716.00A)
16. Pump tubing, 18 (Autoclaveable marprene manifold pump tubing; orange/orange; 0.88 mm bore, Watson-Marlow, 978.0088.00+)
17. Culture chamber jars, 16 (Clear glass powder jars; 500 ml)
18. Culture chamber rubber bungs, 16 (Fisherbrand™ Solid Rubber Stoppers; 45 mm bottom; 51 mm top Fisher Scientific, 41122502)
19. Aeration needle, 16 (Central Surgical Company™ Stainless Steel Needle; 16 G; 203 mm, Fisher Scientific, 12329259)
20. Media influx needle, 48 (B Braun™ Hypodermic Needles Pink 1.2 mm 18 G 50 mm, Fisher Scientific, 10722784)
21. Efflux needle, 16 (Central Surgical Company™ Stainless Steel Needle; 16 G; 101 mm, Fischer Scientific, 12339259)
22. Sampling needle, 16 (Central Surgical Company™ Stainless Steel Needle; 16 G; 101 mm, Fischer Scientific, 12339259 *or* Central Surgical Company™ Stainless Steel Needle; 16 G; 203 mm, Fisher Scientific, 12329259)
23. Male luer cap, 16 (Male Luer Lock Plug; Nylon, Cole Parmer, WZ-45505-56)
24. Collection chamber jars, 32 (Clear glass powder jars; 175 ml)
25. Collection chamber rubber bungs, 16 (Fisherbrand™ Solid Rubber Stoppers; 37 mm bottom; 42.5 mm top, Fisher Scientific, 41122502)
26. 20 L medium container, 2 *optional*
27. Silicone stopper for 20 L medium container, 2 *optional*
28. 5 L medium container, 2 (Pyrex™ Borosilicate Glass Reagent Bottles with Polypropylene Cap and Pouring Ring; 5000 mL, Fisher Scientific, 12094637) *optional*
29. 2 L medium container, 20 (Pyrex™ Borosilicate Glass Reagent Bottles with Polypropylene Cap and Pouring Ring; 2000 mL, Fisher Scientific, 11922629) *optional*
30. Lids with holes for 5 L and 2 L medium containers, 14 (GL45 Screw cap for Pyrex GL 45 media-lab bottle, Fisher Scientific, 15173927) *optional*
31. Syringes, 100 (BD Discardit™ Eccentric Luer-Slip Two-Piece Syringe, Fisher Scientific, 10152534)
32. Linear LEDs (LEDVANCE, 600 10 W, 3000 K warm white)

### PROTOCOLS

#### ASSEMBLING THE MESOSTATS FOR THE FIRST TIME

See item Figure 3–5 for visual representation of how the parts connect. A more detailed description with possible variations in design is given below. Place all vessels and machinery in their intended location before cutting the tubing to ensure sufficient lengths and to minimise mistakes.

- Medium container(s):
  1. Prepare the medium container lids.
    a. If using silicone stoppers:
      i. If the medium container has a tap, the silicone stopper only needs a hole for air to escape. Drill a hole large enough to squeeze medium size tubing through but ensuring a tight seal so that no air can escape around the edges. Attach a short piece of tubing, sticking out about 1 cm on the underside of the stopper and 5 cm on top. Attach a 0.45-micron filter to the tubing.
      ii. If the medium container does not have a tap, a medium siphon is also needed. Drill a second hole in the silicone stopper of the same size and squeeze through medium size tubing long enough to reach the bottom of the container and sticking up about 5 cm on top of the stopper.
    b. If using screwtop lids with holes, first insert a short piece of medium size tubing through one hole so that it ends just inside the lid and sticks out about 5 cm on top. Attach a 0.45-micron filter to the tubing. For the medium siphon, insert a piece of medium size tubing, long enough to reach the bottom of the container and stick out about 5 cm on top of the lid.
  2. Attach the lid or stopper to the medium container.
    a. If there is a medium siphon, attach the long piece of tubing to a reducing connector.
    b. If there is no medium siphon and the medium container has a tap, attach a short piece of appropriate size tubing to the tap and attach this to a reducing connector.
- Media lines:
  3. Attach a short piece of small size tubing to the other side of the reducing connector and attach it to a Y-connector. Split the medium line into as many channels as needed by connecting short (2—3 cm) pieces of small tubing and Y-connectors. Channels receiving the same treatment should share a medium source.
  4. For each channel, attach small size tubing of sufficient length to reach the pump and end with a male luer lock.
  5. For each channel, cut off two 18G hypodermic needles to 0.5 cm and insert them into the ends of a piece of pump tubing. Be careful to not cut up the inside of the pump tubing, if pieces detach, they will cause blockages. The ends of the needles may need additional blunting or filing to reduce sharpness. A damaged piece of pump tubing can easily be cut off.
  6. Connect the pump tubing to the media lines by screwing the needle luer end to the male luer lock. Screw another male luer lock to the other end of the pump tubing.
  7. Cut a piece of small size tubing long enough to reach from the pump to the culture chamber with some slack. Plan carefully where each chamber is going to sit, if multiple treatments are used they should be distributed randomly throughout the array to avoid effects of e.g. differing light level. Attach this piece of tubing to the male luer lock at the end of the pump tubing, and end with another male luer lock.
  8. Label the tubing with autoclave tape so you know which chamber it should connect to.
- Culture and overflow chambers:
  9. For each chamber, prepare the culture chamber rubber bung by drilling four 1 mm holes, one for each hypodermic needle. This aids pushing the needles through, but the seal around them should still be very tight.
  10. Push through one 18 G short needle (medium influx), one 18 G 20 cm needle (air influx) and two 18 G 10 cm needles (culture efflux and sampling needle). A longer sampling needle can be used if desired.
  11. Label the rubber bung so you know which chamber it belongs to.
  12. Mark the 500 ml glass jar at the desired volume (380 ml) as well as ±19 ml (±5%) to make the magnitude of any volume inconsistencies easier to judge by eye.
  13. Attach the media line to the medium influx needle.
  14. Cover the sampling needle with a male luer cap.
  15. Attach a male luer lock to the culture efflux needle and set the needle at the height required for the desired culture volume, i.e. it should be skimming the surface of the culture.
  16. For each culture chamber, prepare an overflow chamber. For each overflow chamber, prepare a rubber bung by drilling two holes large enough to squeeze the lock end of female luer locks into (approximately 2.5 mm in diameter). Connect one hole with a small size of tubing to the male luer lock on the culture efflux needle, and attach another female luer lock on the underside of the bung as a funnel. Attach a short piece of small size tubing to the other hole and end with a 0.2-micron filter.
  17. Attach a male luer lock and push the air influx needle down to touch the bottom of the chamber.
  18. Mount lights around the chambers and/or place the chambers on a light table, using a light meter to ensure light levels are even.
- Aeration system:
  19. Mount the 4-port manifold(s) above the chambers using clamps or tape so that the aeration tubing will be held up without kinks.
  20. Attach a short piece of small size tubing and a 0.2-micron filter to each active port on the manifold.
  21. Connect a small size tube to the air influx needle long enough to reach the 4-port manifold and attach to the other side of the filter.
  22. Prepare the gas washing bottle rubber bung by drilling a hole large enough to push the long glass pipette through, but tight enough that no air can escape around it. Push the pipette through down to the bottom of the flask. If needed, use a sealant around edges of the hole.
  23. Connect the top of the glass pipette to the air supply (building supply or an aquarium pump) with appropriate size tubing.
  24. Put dH2O in the gas washing bottle. The water level should be so that the water sufficiently covers the air outflow from the pipette, but not so that water enters the sidearm when the air is on. Mark upper and lower water levels on the gas washing bottle.
  25. Connect the sidearm to a 0.2-micron filter with a short piece of medium size tubing and the filter to the manifold with a longer piece of medium size tubing. A clamp can be placed on this part of the tubing to control air flow if needed.

#### AUTOCLAVING

All parts that will come into contact with the medium need to be sterilised before use:

1. Medium container(s):
  a. Disconnect the reducing connector on media line between the medium container and the pump tubing.
  b. Prepare the medium and place in the container as it should be autoclaved with the medium inside.
  c. Place a filter in the open tubing on the medium container.
  d. Seal tightly with autoclave bags and autoclave tape around all filters.
  e. Autoclave at 121 (15psi) for 30 min (longer might be necessary for larger containers).
2. Media line:
  a. Disconnect at male luer locks to media influx needle.
  b. Neatly roll up each media line and secure with autoclave tape.
  c. Place all pieces of tubing in an autoclave bag. Autoclave at 121 (15 psi) for 30 min.
3. Culture chamber and collection chamber bungs:
  a. Disconnect filters from tubing to 4-port manifold, and disconnect the male luer locks to the efflux line.
  b. Place the rubber bungs with the needles and tubing in place into autoclaveable trays.
  c. Place the trays in autoclave bags and seal with autoclave tape, taking care to not let the needles pierce the bag.
  d. Autoclave at 121 (15 psi) for 30 min.
4. Culture chamber and collection chamber jars:
  a. These jars are not autoclaveable. Sterilise using 70% IMS and rinse with dH2O.

#### PREPARING FOR AN EXPERIMENT

1. Reconnect all parts after sterilising. Wear gloves washed with 70% IMS at all times, and wear eye protection when handling the hypodermic needles.
2. When the array is assembled, start the pump to fill up the chambers and turn on the aeration system. At max speed (90 RPM) the flow rate is 2.92 ml/minute, meaning it will take approximately 2 hours and 10 minutes to fill the chambers to 380 ml.
3. When the chambers are full, ensure medium levels are equal and set pump to experiment speed. Adjust the efflux needle as necessary and monitor the overflow bottles to ensure efflux is equal.
4. Turn on the lights and ensure control conditions for light level, internal culture and ambient temperatures are met (see Table 1).

#### INOCULATING WITH ALGAE

Inoculating with *C. reinhardtii* from static stock culture:

1. Fill up a couple of 15 ml falcon tubes with stock solution and centrifuge at 2000 RPM for approximately 20 minutes. If there is a red layer on top of the green pellet, remove it as this is bacterial contamination. Pour out the supernatant and mix the pellet with fresh, sterile medium. Repeat washing procedure twice.
2. Use a sterile syringe to push an equal amount of freshly washed algal cells into each chamber through the sampling needle. Note: The inoculation volume will vary depending on the amount of stock algae available, the stock density and the desired starting density for the cultures.
3. Allow cultures to reach steady state before applying experimental treatments. See below for methods for how to sample to estimate concentration.

#### APPLYING TREATMENTS

Treatments can be applied gradually through the medium line by adding the compound directly to the medium or as shock injections through the sampling needle. The two methods can also be combined. For shock injection:

1. Prepare a mixture of medium and compound at as high a concentration as practical to ensure the volume injected into the chamber is as small as possible and reducing the effects of dilution of the culture.
2. Before injection, remove culture at a volume corresponding to the intended injection volume by using a sterile syringe to pull it out through the sampling needle. This so the treatment can be adequately mixed into the culture, and to avoid a sudden rush of culture through the efflux line.
3. Inject the treatment in through the sampling needle using a sterile syringe. All chambers should receive the same injection volume, including controls.

#### DAILY MAINTENANCE

To ensure equal conditions in all chambers, the mesostat array must be attended to daily according to the daily maintenance protocol:

1. Check water level in gas washing bottle is between max and min markings. Top up with distilled water if running low. Avoid going over the max marking, as this will cause water to bubble into the airflow tubing which can lead to blocked filters.
2. Check culture chamber airflow is satisfactory and equal. If a culture chamber has low airflow, check if the filters are blocked, first the filter on the adjoining collection chamber, then the filter connecting to the manifold, and replace if necessary. The most common cause of filter blockage is them becoming wet, either by an efflux blockage or low pressure resulting in the culture entering the aeration needling and tubing, or by high ambient humidity. Make a note of airflow problems data is being collected if it is possible the problems have been present for more than an hour. If filters are often blocked, consider changing the ambient humidity.
3. Check for leaks around connectors and luer locks. If leaking, first try tightening. If that does not help, replace with a new part (sometimes autoclaving can warp the luer locks), taking care to sterilise the new part with 70% IMS.
4. Check culture levels are even and do not deviate from the 380 ml line. If the medium level is too low, check the media influx tubing and needle for blockages. The most likely points for blockages are inside the pump tubing and any needles due to their narrow gauges. If medium influx is normal, adjust the efflux needle, and check if the level is back to normal in a couple of hours (time needed to wait dependent on flow rate and total volume deviation). If the level is too high, examine the efflux tubing and needle for blockages, along with the collection chamber filter. When unblocked, the culture chamber level should return to normal volume relatively quickly. Always make a note of culture level changes if data is being collected.
5. Check collection bottle levels are even, and measure volume of a few to ensure the flow rate is correct. Empty the collection bottles and note the time, so that flow rate can be calculated when the bottles are next emptied.

#### SAMPLING AND MONITORING

1. Temporarily restrict the airflow using the adjustable clamp on the tubing. This prevents liquid from bubbling up through the sampling needle.
2. Wearing gloves washed with 70% IMS, unscrew the cap on the sampling needle and use a sterile syringe to extract liquid.
3. Put the sample in a labelled Eppendorf tube and put the cap back on the sampling needle, ensuring it is screwed on tightly. Clean up any spillage.
4. Repeat for each chamber.
5. Do not forget to turn the airflow back on.
6. The samples may be stored for later processing or counting, either through flash freezing with LN_2_ and subsequent storage at −80°C, or by mixing with Lugol’s solution and storing in a fridge at 4°C depending on the intended use for the samples.

## ASSEMBLY SCHEMATIC

### EXPERIMENTAL DESIGN FOR VALIDATION DATA

#### REPLICABILITY

Presented below are control data from four separate experiments using the linked protocol to show replicability. The conditions and relevant differences for these experiments are summarised in Table 2, unless otherwise stated the experimental conditions correspond to those outlined in the protocol. In all of the presented experiments, *Chlamydomonas reinhardtii* strain Sager’s CC-1690 wild-type 21 gr was used, obtained from the *Chlamydomonas* Resource Centre (University of Minnesota, St Paul, MN, USA) core collection. Two different dilution rates were used in the experiments: 0.3/day and 0.15/day. The former was based on the dilution rates used in previous experiments using chemostat populations of similar species that this system was designed for (e.g. Becks *et al*., 2012; Yoshida *et al*., 2003), the latter was used as an alternative lower rate to decrease the consumption of growth medium as well as wear and tear on the pump tubing.

**Table 2:**
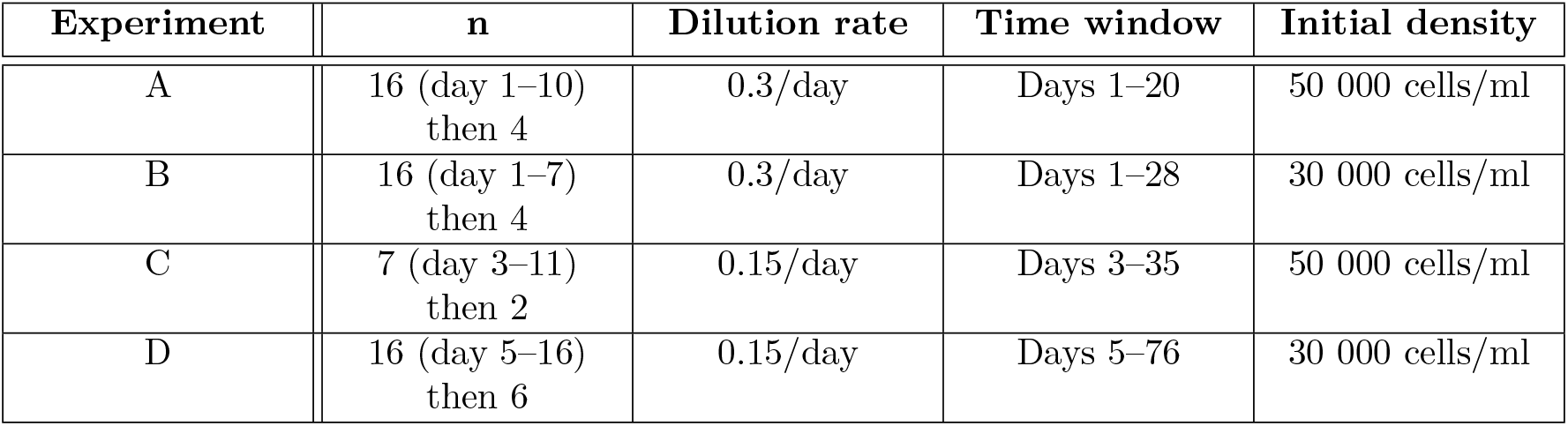
Summary of experimental conditions and properties of data used in Figure 6–10. Data from all four experiments were used to generate Figure 6, whereas data from experiment C was used for Figure 7, and experiment D data were used for Figure 8, Figure 9 and Figure 10

| Experiment | n | Dilution rate | Time window | Initial density |
| --- | --- | --- | --- | --- |
| A | 16 (day 1–10)<br>then 4 | 0.3/day | Days 1–20 | 50 000 cells/ml |
| B | 16 (day 1–7)<br>then 4 | 0.3/day | Days 1–28 | 30 000 cells/ml |
| C | 7 (day 3–11)<br>then 2 | 0.15/day | Days 3–35 | 50 000 cells/ml |
| D | 16 (day 5–16)<br>then 6 | 0.15/day | Days 5–76 | 30 000 cells/ml |

#### APPLICABILITY TO EXPERIMENTAL HERBICIDE RESISTANCE EVOLUTION

Six mesostat chambers in experiment C were allowed a week to reach steady state before the glyphosate treatment was introduced. Shock injections of 38 ml were performed as described in the protocol bringing two chambers each to concentrations of 0 mg/L (controls), 100 mg/L and 150 mg/L glyphosate (analytical standard, PESTANAL®). Both of the chosen glyphosate concentrations are above the minimum inhibitory concentration for *C. reinhardtii* of 97.5 mg/L (Lagator *et al*., 2013b).

#### COMMON PROBLEMS

We have provided data from three common problems that present with this type of system: a leak, contamination and algal clumping, all from experiment D. These were spontaneous events and the data presented here aims to show how to identify their signal in the population density data and distinguish it from normal variation among populations. The leak in this example resulted in elevated dilution of a single chamber for roughly four hours due to a clamp securing the pump tubing cassette coming undone. In the case of the contamination event, all of the presented six chambers had been disconnected from the array six days before bacterial contamination was observed under the microscope in four chambers, with the remaining two unaffected by the contamination event. The clumping phenotype was not receiving control medium but presented in a population undergoing treatment with a sublethal dose of glyphosate.

### SAMPLE PROCESSING

Population density was in all cases determined through flow cytometry (Beckman Coulter CytoFLEX), using CytExpert (Beckman Coulter) to gate and count events detected in the PerCP-A channel (Excitation: 488nm, Emission: 690/50 BP). This channel is used to detect chlorophyll *a* and represents a robust method for estimating algal density (Kadono *et al*., 2004) which was further validated against manual haemocytometer counts for this system.

### DATA HANDLING

All statistical analyses were carried out in R (version 4.0.5, (R Core Team, 2021)), using the `lme4` package (Bates *et al*., 2015) to fit a linear mixed effects model with log-transformed population density as the response, dilution rate and experiment as fixed effects, and day and chamber as random effects with varying intercepts. The significance of the fixed effects was tested using the Anova() function from the `car` package (Fox & Weisberg, 2019) and confirmed through parametric bootstrapping using the `pbkrtest` package (Halekoh & Højsgaard, 2014).

The slope of population density decline was estimated between days 6–16 with the package `emmeans` (Lenth, 2022) after fitting a linear mixed effects model with the log-transformed population density as the response, treatment and day as fixed effects as well as day and chamber as random effects with varying intercepts.

## RESULTS

### REPLICABILITY

Across the experiments presented here, there is no difference between the mean population densities after steady state has been reached (*χ*^2^ = 2.1, DF = 3, p = 0.6, Figure 6). Furthermore, there was no difference in steady state population density whether the dilution rate is 0.3 or 0.15/day (*χ*^2^ = 0.4, DF = 1 p = 0.5). The length of the establishment batch phase before steady state is reached will differ depending on the conditions and inoculate density. The dynamics during this phase has the potential to affect the makeup of the population and thus later dynamics, and it is thus advisable to let the cultures reach steady state before introducing treatments. However, all experiments presented here reached steady state within the first week and it was maintainable for several weeks thereafter.

**Figure 6:**
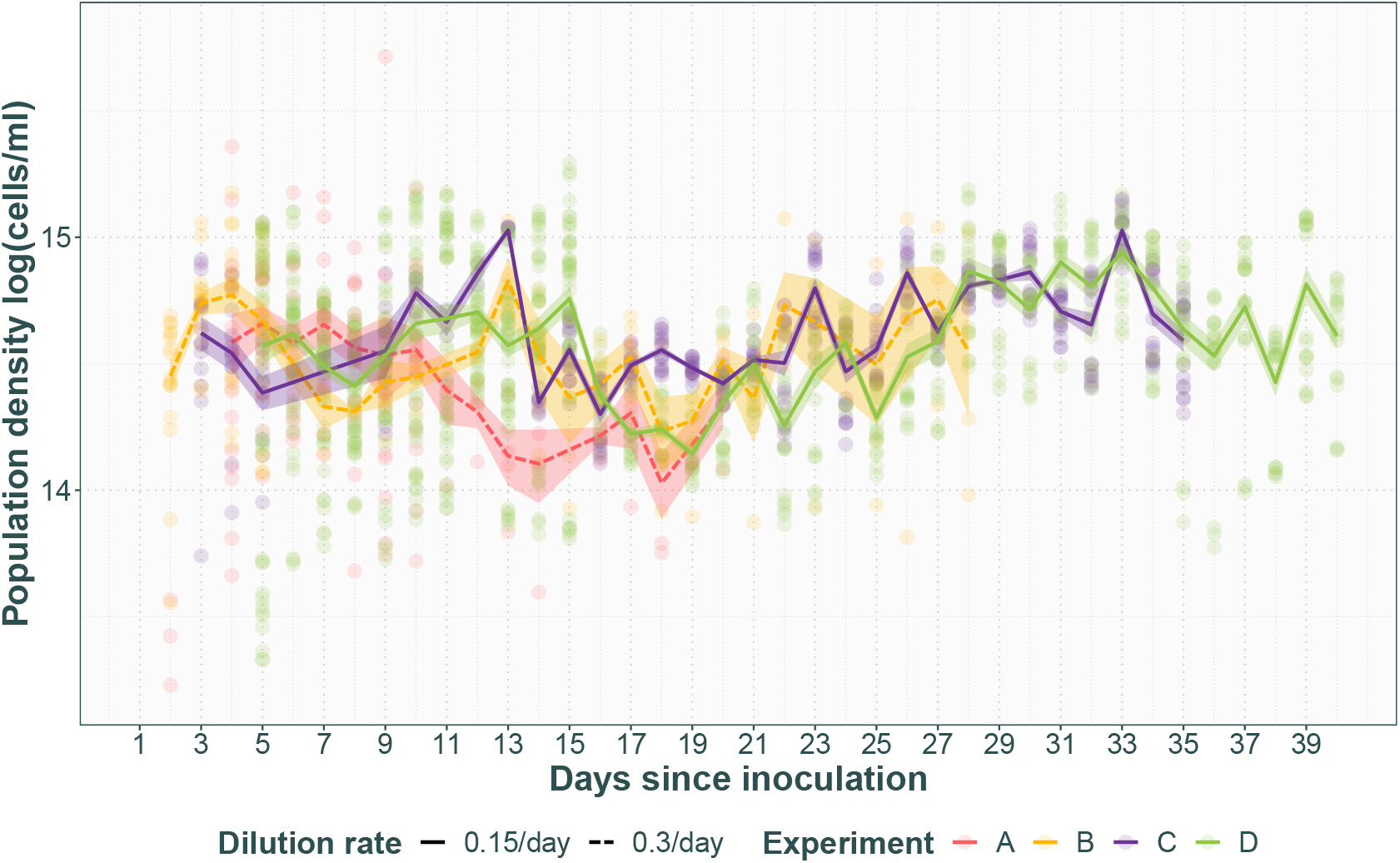
Population density with time in four separate runs of the mesostat system. Transparent points represent technical replicates and opaque lines with standard error represent average across populations for experiment. Experiments A and B had a dilution rate of 0.3/day (dashed line), whereas experiments C and D had a dilution rate of 0.15/day (solid line). Note that all have runs have a brief and rapid decline in population density between day 11 and 16. This corresponds to an injection of additional medium as part of the experiment the data is from.

The level of variation observed in this data set is normal for this type of system (Becks *et al*., 2012; Fussmann *et al*., 2000; Yoshida *et al*., 2003) and can be divided into among population variation and day-to-day variation. Among population variation is primarily caused by the biology of the system as these are separate, genetically heterogeneous populations on separate evolutionary trajectories. Day-to-day variation is however at least partly caused by limitations in sample processing. Both are discussed in more detail in the Discussion section, as well as how to reduce or circumvent the latter in particular.

### APPLICABILITY TO EXPERIMENTAL HERBICIDE RESISTANCE EVOLUTION

Figure 7 shows the population densities of the four glyphosate treated populations and two control populations for 24 days following glyphosate treatment introduction. The glyphosate treated chambers exhibit population decline at a rate approximate to (150 mg/L, slope = −0.14, SE = 0.006) or below (100 mg/L, slope = −0.098, SE = 0.006) the dilution rate of 0.15/day. In the same timespan, the control populations exhibit an overall slight increase in population density (slope = 0.022, SE = 0.006), possibly reflecting adaptation to the mesostat environment. The onset of the population decline appears to be immediate for the 150 mg/L glyphosate treatment, whereas it occurs roughly 5 days after the glyphosate injection for the 100 mg/L glyphosate treatment. This is likely due to the 100 mg/L glyphosate treatment being just on the cusp of the minimum inhibitory concentration, enabling the populations to maintain growth for a short while before the herbicidal action is apparent. After 15 and 18 days respectively of population density decline, the 100 mg/L populations increase in cell density again, suggesting the populations have evolved resistance to the glyphosate, whereas the 150 mg/L populations never show evidence of resistance.

**Figure 7:**
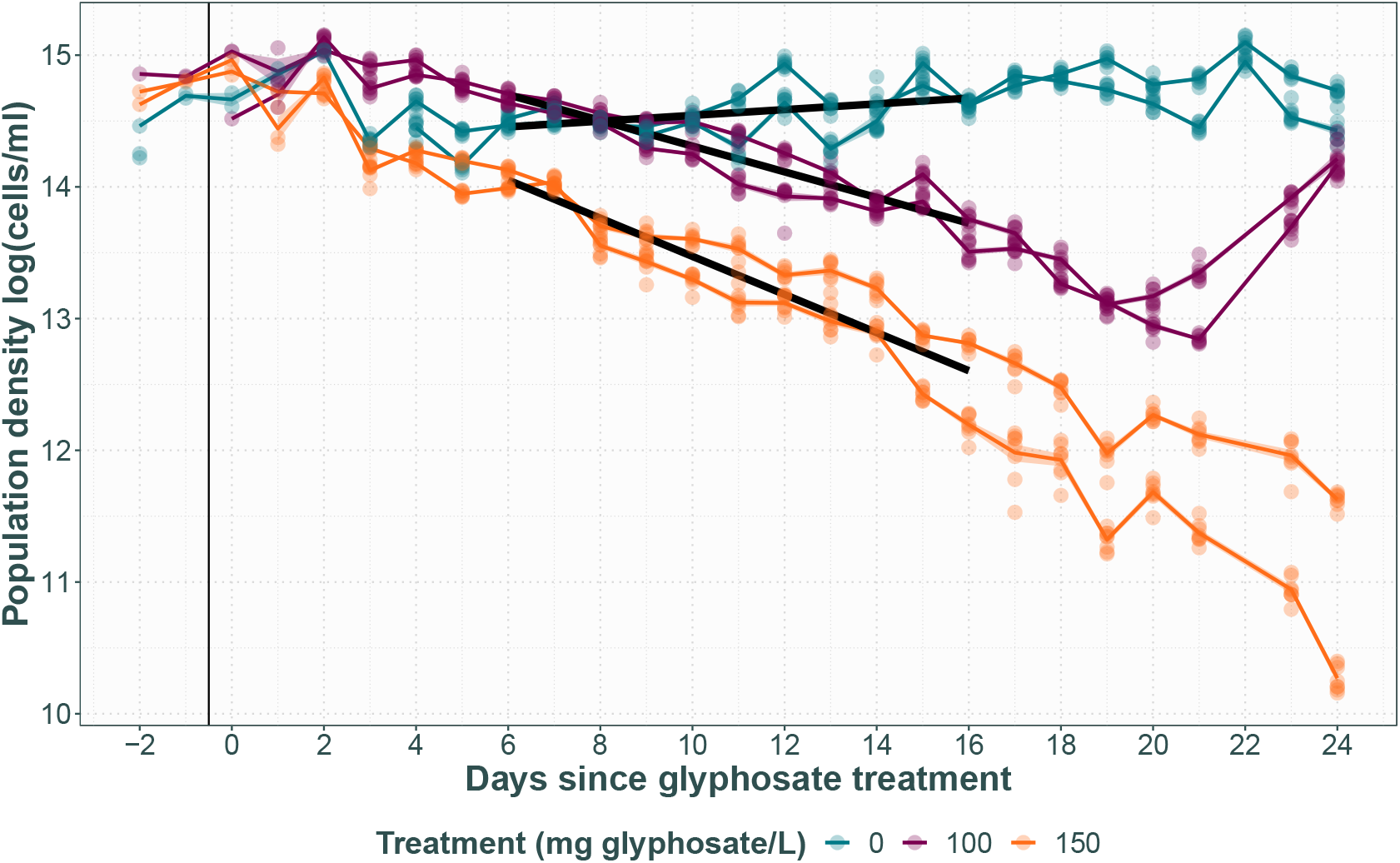
Population density with time in populations receiving 0, 100 or 150 mg/L glyphosate. Transparent points represent technical replicates, with opaque lines for population average with standard error transparent ribbon. Thick black lines represent the fitted linear model and the thin black vertical line shows start of the treatment.

### COMMON PROBLEMS

#### LEAKS

Figure 8 shows the population density in chamber F after a major leak causing over-dilution. Compared to the expected among population and within-population day-to-day variation observed in the chambers that did not experience a leak, three crucial differences together make this the characteristic signal of over-dilution: 1) While similarly large day-to-day fluctuations in the measured density occur in the presented data set, day effects present across chambers. The rapid reduction in population density for chamber F between days 34 and 35 is only apparent in that chamber, whereas a similar reduction between days 37 and 38 is seen in all of chambers A–E. 2) The reduction in population density in chamber F results in a lower population density than otherwise observed in the data set (by roughly 3 × 10^5^ cells/ml). 3) The reduced population density is observed in chamber F for several days after day 35, rather than recovering by the next day like seen for chambers A–E after day 38.

**Figure 8:**
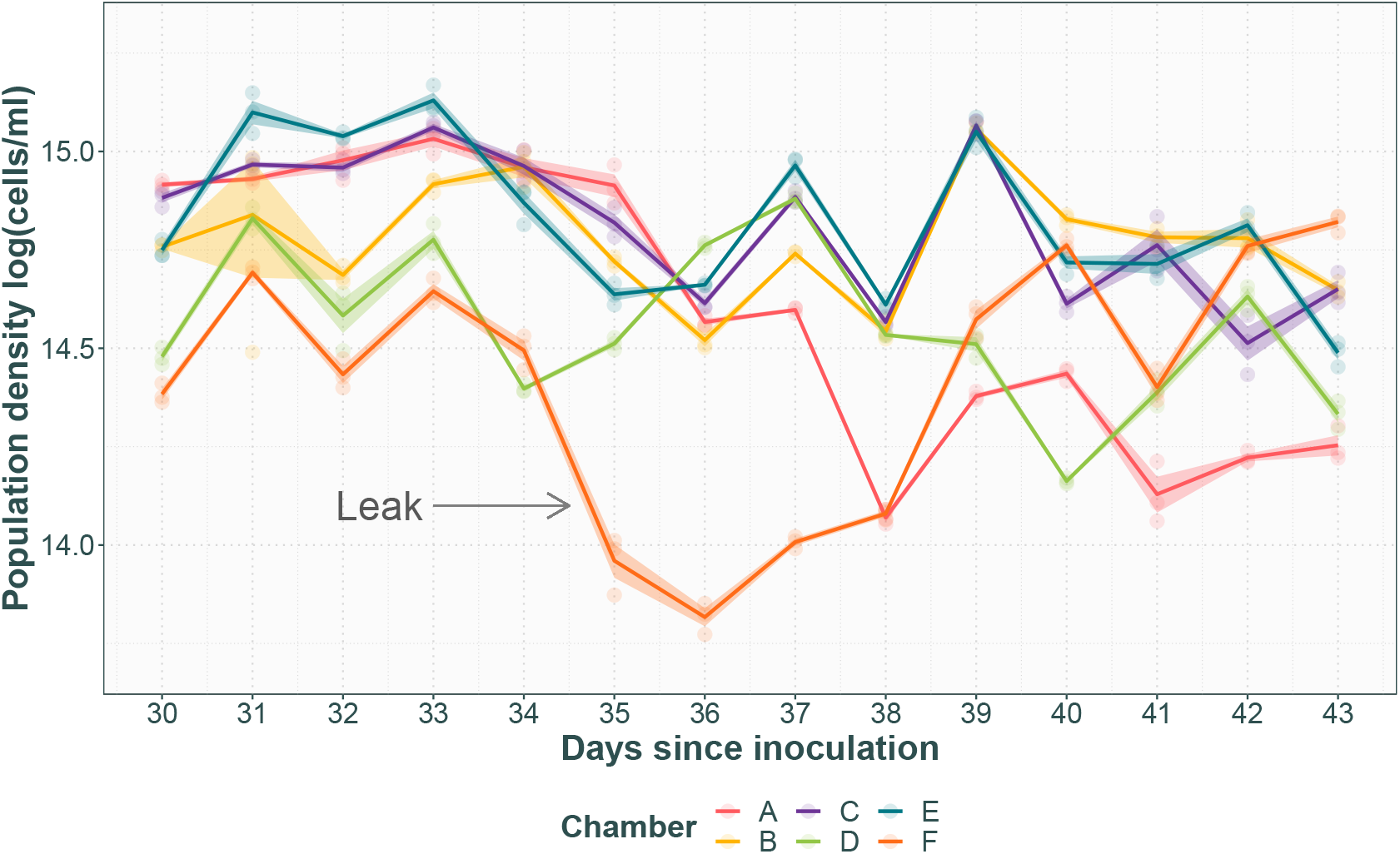
Population density with time after a major leak. Transparent points represent technical replicates with opaque lines for population average with standard error. The leak caused overdilution of chamber F between days 34 and 35 (indicated by arrow), compared to the unaffected chambers A—E.

#### CONTAMINATION

Figure 9 shows the gradual population density decline in four chambers where bacterial contamination was observed under the microscope compared to two chambers that were unaffected by the contamination event. While the average population density of the contaminated chambers starts to trend lower a few days after the contamination event, the full effect on the population density is not clear until several days after the contamination had been observed under the microscope. Furthermore, while there is considerable variation among all populations, the signal of contamination in the data is clearly distinguished from the expected among population variation and day-to-day variation by the fact that it is a consistent, long-term population-density decline without recovery 12 days after the contamination event.

**Figure 9:**
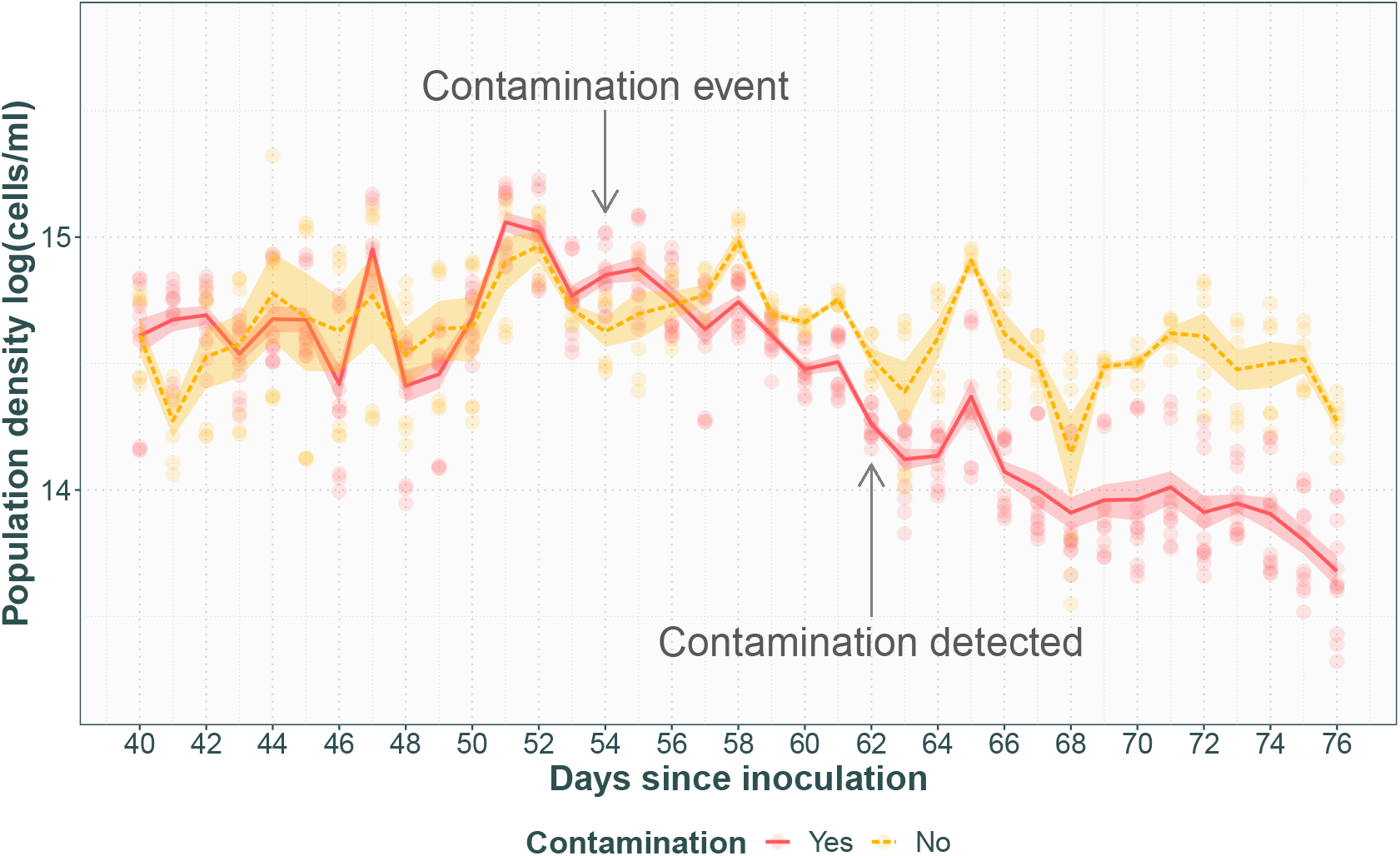
Population density with time after a contamination event. Transparent points represent technical replicates with opaque lines with standard error for average of contaminated (solid line) vs. non-contaminated (dashed line) populations. Contamination is likely to have entered the system at day 54 (indicated by arrow), and bacterial contamination was found in 4 out of 6 chambers on day 62 (indicated by arrow).

#### CLUMPING

Figure 10 shows flow cytometry population density estimates from a population exhibiting a clumping phenotype compared to non-clumping populations undergoing the same treatment. The data signal here is an artefact of the limitations of the instrument being unable to accurately distinguish individual cells within aggregates, resulting in huge fluctuations in estimated cell density considerably larger than and out of step with the day-to-day variation observed in the other populations.

**Figure 10:**
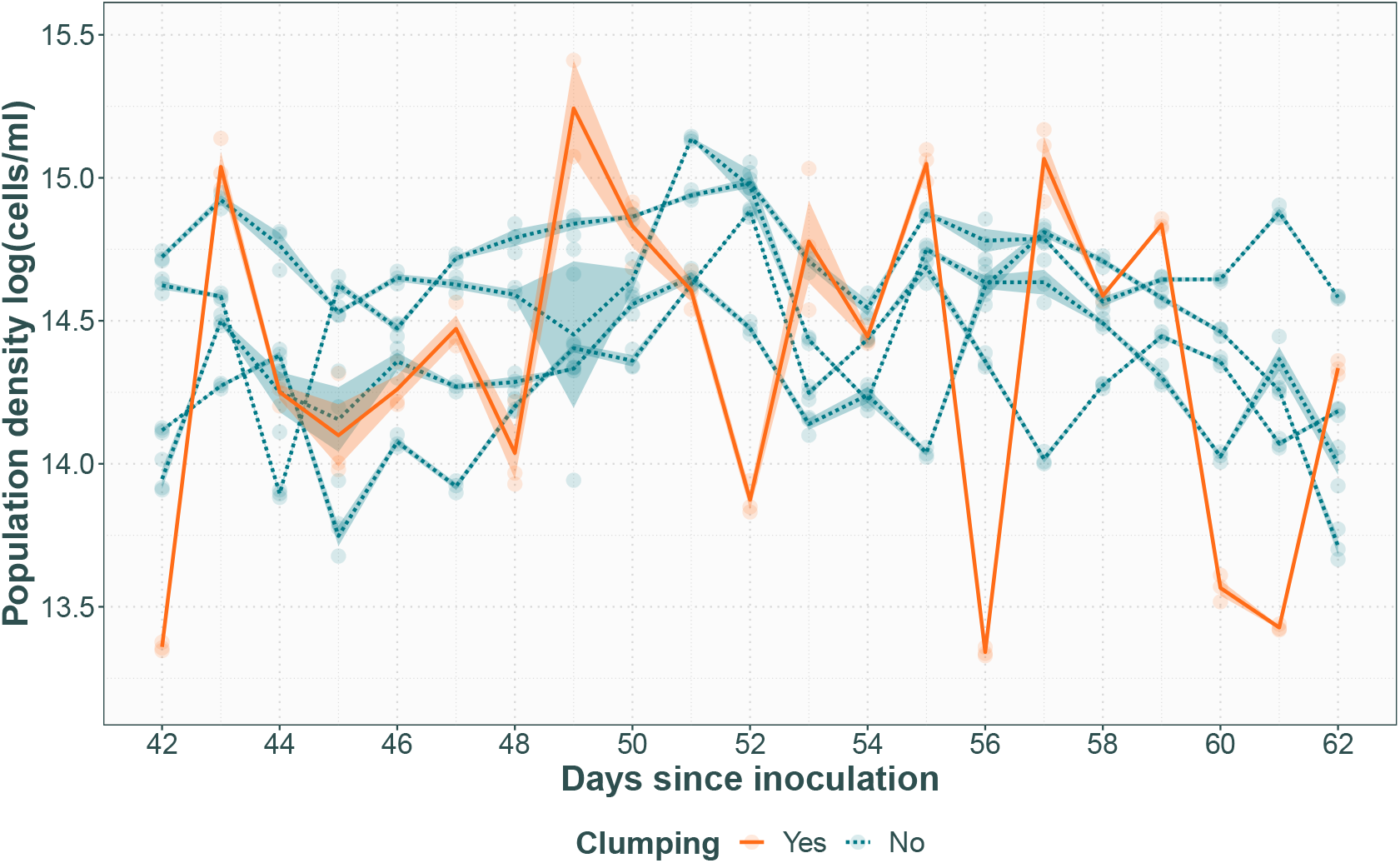
Population density with time in a population exhibiting a clumping phenotype. Clumping population shown with solid line, compared to four other populations receiving the same treatment that did not exhibit clumping in dashed line. Transparent points represent technical replicates, with the lines for population averages with standard error.

## DISCUSSION

Chemostats offer a number of advantages over batch cultures for long-term experimental evolution research. Precise control of selective pressures in a chemically constant environment without evolutionary bottlenecks along with a link between growth rate and dilution rate constitute a useful conceptual framework for modelling evolutionary adaptation and population dynamics. This system adds to the small, but growing, number of efforts to produce simple but scalable, multiplexed DIY chemostats from cheaper materials that are possible to build and maintain by a single person (Dénervaud *et al*., 2013; Ekkers *et al*., 2020; Miller *et al*., 2013; Skelding *et al*., 2018; Tonoyan *et al*., 2020; Toprak *et al*., 2013; Wong *et al*., 2018), and is the first of its kind for experimental evolution of algae, specifically the evolution of herbicide resistance in model species *C. reinhardtii*. There are three substantive changes from the Miller *et al*. (2013) ministats, one system specific and two generic changes to suit experimental evolution with continuous sample extraction. Firstly the system was adapted to suit the study species *C. reinhardtii*, including light and a lower dilution rate, which distinguishes the system from previous DIY chemostat arrays developed for maintenance of yeast (Dénervaud *et al*., 2013; Miller *et al*., 2013; Wong *et al*., 2018) and bacterial cultures (Tonoyan *et al*., 2020; Toprak *et al*., 2013). Secondly, a needle and syringe system was added to facilitate easy, sterile access to the culture for the removal of samples. This allows sampling from the middle of the active culture rather than relying on the overflow. The efflux only samples from the top and the overflow chamber constitutes a wholly different environment without continuous dilution, build-up of waste products and increased evaporation, making them unrepresentative samples of the chamber populations. Furthermore this simplifies addition of cells or treatment compounds directly to the chambers, eliminating the risk of contamination that comes with disconnecting the medium influx or efflux channels. While sampling ports have been described before (e.g. Ekkers *et al*., 2020; Tonoyan *et al*., 2020) our simplification and combination with syringe extraction allows manual sampling with minimal contamination. The third change is an increase in the chamber volume to allow larger population sizes and possible future introduction of several trophic levels. Furthermore, this increases increases the amount of sample that can be extracted on a regular basis, extending the possibilities for the types of assays that can be performed to characterise evolution in action, as most of the previous DIY chemostat arrays have been limited by their small working sizes (Ekkers *et al*., 2020; Miller *et al*., 2013; Tonoyan *et al*., 2020). Lastly, there were several changes to specific materials to lower the overall costs.

### SOURCES OF VARIATION AND HOW TO MINIMISE IT

The data presented here illustrates the expected variation between cultures and how to identify the signal of equipment failure, such as a leak, or contamination. We also demonstrate that the system can be used to evolve resistance to growth inhibiting herbicide glyphosate, and that the signal of herbicidal action is apparent as a population density decline, followed by an increase after the population has evolved resistance. The herbicidal effect is clearly distinguishable from the expected variation under control conditions, and given enough time, the resistant population is expected to settle at a new steady state.

The variation among replicate populations observed here is normal (Becks *et al*., 2012; Fussmann *et al*., 2000; Yoshida *et al*., 2003) and expected as they constitute separate, genetically heterogeneous populations on separate evolutionary trajectories. Even when using a single founder population, the genetic bottleneck caused by splitting it between populations as well as the dynamics during the establishment phase of batch-like growth dynamics (Gresham *et al*., 2008) will result in similar but distinct populations by the time they reach steady state. Effort should be made to ensure that all chambers receive the same levels of light and aeration as well as consistent dilution with the same medium, and starting variation could be eliminated through starting with clone populations at a high enough concentration to effectively avoid the establishment phase. However, the among population variation is generally of scientific interest to experimental evolution studies and should be investigated rather than eliminated.

Conversely, while day-to-day variation within a population is also normal for this type of system, it is also partly caused by limitations to the sampling protocol. The data presented here was obtained from measurements performed on living cells that had the opportunity to grow and divide between sample extraction from the mesostat chambers and sampling processing. While this is an unavoidable source of variation, it can however be reduced by minimising the time that passes and working in a controlled temperature environment. If the experimental design allows, the cells can be immobilised by using e.g. Lugol’s solution before counting with flow- or haemocytometry. It is also possible to control for this variation by including sampling day as a source of error in statistical models applied to the data.

The among population and day-to-day within population variation are however both clearly distinguishable from the data signal of common faults like leaks, contamination and clumping. While these faults are likely to be detected before they become apparent in the population density data, leaks causing significant over-dilution are apparent within a few hours while clumping and contamination can be observed under a microscope, it is important to understand how they affect the data so that an informed decision can be made on how to handle it. While the population density is always expected to quickly return to steady state after over-dilution, the increased flushing out of cells constitutes an evolutionary bottleneck and the changed growth conditions may affect other traits of the population not visible in the population density data and data collected subsequent to a major leak should thus be treated with caution. The leak presented here was caused by equipment failure resulting in over-dilution, but smaller leaks often occur as the pump tubing wears out with long term use, which can lead to under-dilution of the connected chambers. Both are best prevented by regular inspection of the pump parts for irregularities.

Bacterial contamination is another common risk in long-running continuous cultures (Gresham & Hong, 2014), and is best prevented by working in a sterile environment and minimising the points at which contamination can enter the system. The main contamination risk presents when disconnecting any part of the array, such as when switching medium containers, or when extracting samples, and particular care should be taken to keep the connecting parts sterilised during. The example presented here is the only instance of contamination observed across eight separate experiments each lasting more than a month and happened when the chambers were disconnected from the array for a longer period of time and removed from the sterile environment. Even so, only four out of six chambers showed evidence of contamination under the microscope 12 days after the contamination event, despite all of the chambers in question sharing a medium source. This suggests that the system is robust in terms of contamination not spreading between the chambers. While regular microscopy inspection of cell samples for contamination is recommended, this can be laborious with a large number of replicates and the characteristic population density decline provides another opportunity to detect and isolate the problem.

Lastly, chemostat populations being under a selective pressure to evolve phenotypes reducing their risk of being flushed out is an often cited issue with the method (Gresham & Dunham, 2014; Gresham & Hong, 2014), presenting as adhesion to the chamber walls and cell flocculations. While this phenomenon has as of yet never been observed under control conditions with this protocol, there was one instance of cells exhibiting a clumping phenotype while under treatment with a sublethal dose of glyphosate, making it possible it was a response to the treatment rather than the mesostat environment. In *C. reinhardtii* there are two distinct types of clumping: cell aggregations of separate cells and palmelloid colonies that share a cell wall (de Carpentier *et al*., 2019; Harris *et al*., 1989; Lürling & Beekman, 2006). Both have been found to be an induced response as well as heritable (Harris *et al*., 1989; Iwasa & Murakami, 1968; Khona *et al*., 2016; Lürling & Beekman, 2006; Olsen *et al*., 1983), meaning that once they become common in a population they may be hard to get rid of (Harris *et al*., 1989). Palmelloids are small enough that they will not cause blockages, but due to the shared cell wall they cannot be disassociated through bubbling or by vortexing a sample. Cell aggregations can be considerably larger, however they are also possible to break apart through vortexing, and vigorous bubbling of the cultures often prevents their formation (Gresham & Dunham, 2014). How much of a problem clumping is depends on the experiment, i.e. it becomes a problem if it hinders sample processing and when it is thought to be an artefact of the chemostat environment rather than in response to the applied treatment. For population density measurements by flow cytometry as presented here, clumping considerably reduces the accuracy of the measurements as each clump is counted as a single particle, increasing day-to-day variation. In this case, manual haemocytometry could give a better estimate but this is considerably more laborious.

### OTHER POSSIBLE ISSUES AND LIMITATIONS

Despite the many advantages of chemostat cultures, there are limitations to their application and caveats to how the data may be interpreted. While the system described in this protocol was explicitly designed to be maintainable by one person as well as cheaper than the Miller *et al*. (2013) ministats it is based on by choosing alternative materials and using parts not purpose bought for this experiment, it is still considerably more expensive than batch cultures. While it is theoretically possible to run very large cultures indefinitely, the cost of the medium or treatment components will limit the lifespan of the experiment as they will be consumed faster than in a batch culture design. One way to conserve medium and treatment components is to lower the dilution rate, which in the experiments presented here had no effect on population density in the chambers. However, this changes the selection pressures experienced by the populations as well as their doubling rate (Gresham & Hong, 2014). The logistics of the system and any cost saving measures must therefore be carefully balanced against the resulting biology, taking into account the desired selective pressure, cell cycle stage and generation time.

This design introduced sampling needles to allow sampling directly from the culture as an alternative to sampling from the overflow chamber, as the environment therein will be different from the culture chamber, or redirecting the overflow, as the low flow rate made sampling a slow process and the high temperature caused high levels of evaporation. However, sampling directly from the culture does perturb the steady state and change the dynamics within the chamber by temporarily reducing the culture volume and thus pausing dilution (Fischer *et al*., 2014). The frequency and volume of samples should thus be carefully considered against the disruption they may cause.

Another potential problem involves insufficient aeration or efflux blockages causing over- or underpressure in the chambers. Provided the air supply is sufficient, the most common reason for low or uneven bubbling is blocked air filters, usually because they have become damp. If the air filters frequently become damp, the ambient humidity may be too high. Not enough bubbling may cause sedimentation and stratification of the culture, as well as selection for phenotypes that sink so they avoid being flushed out, or it may instead cause the culture to rise through the aeration needle instead of through the efflux needle, changing the effective dilution rate. Clogging of the media line is uncommon, but can occur if not properly sterilised and contamination is allowed to grow. This is often apparent as a reduction in flow into and out of the affected chambers. The daily maintenance part of the protocol outlines how to spot and address these problems.

Lastly, in terms of studying the dynamics of adaptive evolution, chemostat systems are highly specific environments. When transplanted out, chemostat populations are often found to grow poorly in their ancestral environment compared to the ancestral strain (Gresham & Hong, 2014; Wenger *et al*., 2011), as they have had intense selection on a specific part of the growth cycle in an environment of constant dilution that is not reflective of natural populations. However, this is also part of their usefulness and beauty, by keeping the adaptive environment as simple and specific as possible, we can isolate fitness effects and allow fine-tuned investigation of their mechanics and dynamics.

### HOW TO IMPROVE OR MODIFY

Several further modifications are possible for this system. A light table that does not transfer heat to the cultures would allow the internal culture temperature to be set solely by the ambient temperature in the controlled temperature room while maintaining low evaporation. As the chamber lids are relatively large, sensors to monitor e.g. pH or CO_2_ levels could also be fitted through additional ports (see Ekkers *et al*., 2020).

While the pump and pump-tubing are integral to the design and also the most expensive parts, all other parts could be easily substituted depending on availability or cost constraints. The materials list provided in the protocol can be used as a guide for the dimensions and properties of the part, but primarily aims to illustrate how this type of system can be built from parts already found in most wet labs rather than buying a pre-made set. Any water-tight, sterilisable container can be used for culture chambers if suitable lids can be manufactured, such as falcon tubes (Tonoyan *et al*., 2020) or commonly available lab glassware (Ekkers *et al*., 2020). The controlled temperature room can be replaced with water baths (note however that this requires mounting the lights up on the sides of the water baths), and portable aquarium pumps can be used instead of building infrastructure, increasing flexibility in where the system can be housed.

The light system here is rudimentary but sufficient for *C. reinhardtii* growth (Harris *et al*., 1989), using white light LED strip lights mounted around the chambers along with a DIY light box also consisting of white light LED strip lights and a semi-transparent plastic top to diffuse the light. The light box is not necessary, but convenient for maximising light from all angles. Under control conditions 24h light was used, but it is possible to fit a timer to the outlet connecting the lights to instead provide a diurnal light cycle. Coloured semi-transparent plastic could be used to provide light only from a specific part of the light spectrum, but it would also be possible to mount specialist lighting around the mesostats if tuning for a specific photosynthetic organism or experiment is desired.

### FUTURE RESEARCH OPPORTUNITIES

We have used this system for experimental evolution of herbicide resistance in algae by adding glyphosate as a shock injection and then continuously through the growth medium, however, this setup is also easily adaptable depending on the research question. The herbicide treatment could also be applied gradually through the medium or through series of shock injections in a ratchet protocol (Reboud *et al*., 2007) and investigate to what level the resistance can be pushed and at what speed. The dilution rate and thus the cell growth rate is set by the pump speed, tubing thickness and culture volume, so running chambers with different dilution rates simultaneously would be possible with different pump tubing thicknesses, multiple pumps or multiple culture volumes, depending on the range required. Furthermore, the use of multiple light tables with opaque partitions between cultures would allow testing for an interaction with light level, or the chambers could be kept in water baths at different temperatures to determine the effect of temperature.

In addition to testing the effect of abiotic factors such as temperature or light, or manipulating the specific cell cycle stage of the population, a particularly interesting future application would be to use the system to ask focused questions about eco-evolutionary dynamics. In particular, introducing several trophic levels in the culture chambers to study the ecosystem and food web effects of herbicides and evolving herbicide resistance. The predator-prey cycles of rotifer *Brachionus calyciflorus* and *C. reinhardtii* as well as *Chlorella vulgaris* have been successfully studied and modelled using chemostat environments (e.g. Becks *et al*., 2012; Yoshida *et al*., 2003) and our setup allows simplified simultaneous replication for this type of system that can be maintained by one person. Competition could also be introduced to the system through using multiple algal strains and monitoring their frequencies or through expanding the culture ecosystem to include other algal species or bacteria (Raatz *et al*., 2018).

## ACKNOWLEDGEMENTS

Special thanks to Allison Blake and Lynsey Gregory for technical support in the lab.

## RIGHTS RETENTION AND DATA AVAILABILITY

For the purpose of open access, the author has applied a Creative Commons Attribution (CC BY) licence to any Author Accepted Manuscript version arising. The data presented in this manuscript, along with relevant code, is available at DOI: 10.5281/zenodo.6786427

